# Differences in high-definition transcranial direct current stimulation over the motor hotspot versus the premotor cortex on motor network excitability

**DOI:** 10.1101/487488

**Authors:** Stephanie Lefebvre, Kay Jann, Allie Schmiesing, Kaori Ito, Mayank Jog, Nicolas Schweighofer, Danny JJ Wang, Sook-Lei Liew

## Abstract

The effectiveness of transcranial direct current stimulation (tDCS) placed over the motor hotspot (thought to represent the primary motor cortex (M1)) to modulate motor network excitability is highly variable. The premotor cortex—particularly the dorsal premotor cortex (PMd)—may be a promising alternative target to more effectively modulate motor excitability, as it influences motor control across multiple pathways, one independent of M1 and one with direct, modulating connections to M1. This double-blind, placebo-controlled study aimed to differentially excite motor and premotor regions using high-definition tDCS (HD-tDCS) with concurrent functional magnetic resonance imaging (fMRI). HD-tDCS applied over either the motor hotspot or the premotor cortex demonstrated high inter-individual variability in changes on cortical motor excitability. However, HD-tDCS over the premotor cortex led to a higher number of responders and greater changes in local fMRI-based complexity than HD-tDCS over the motor hotspot. Furthermore, an analysis of individual motor hotspot anatomical locations revealed that, in more than half of the participants, the motor hotspot is not located over anatomical M1 boundaries, despite using a canonical definition of the motor hotspot. This heterogeneity in stimulation site may contribute to the variability of tDCS results. Altogether, these findings provide new considerations to enhance tDCS reliability.

## INTRODUCTION

Transcranial direct current stimulation (tDCS) is a noninvasive tool that can modulate cortical excitability in human brain regions^1^. The effects of tDCS on upper limb motor control and motor rehabilitation have been widely studied (for a review see Lefebvre & Liew^2^), and promising effects of tDCS applied over the primary motor cortex (M1), functionally localized as the motor hotspot^3 4,5^ in each participant, have been demonstrated in post-stroke motor recovery^6-10^. However, a major issue that prevents tDCS from being widely adopted in clinical practice is the high inter-individual variability shown in motor behavioral or cortical physiological changes following M1 stimulation^11,12^. The reasons for this inter-individual variability have not been fully elucidated^13^.

One potential solution to reduce the inter-individual variability is to use alternative stimulation sites and/or more precise tDCS montages. In particular, stimulating another entry point into the motor network may lead to more reliable changes following tDCS. The premotor cortex, and particularly the dorsal premotor cortex (PMd), is a promising alternative neural target for modulating motor excitability as it is the second major origin of the corticospinal tract (CST), with 30% of CST projections^14,15^. Non-human primates studies have shown that PMd provides high-level control of the upper limb motor commands using a downstream pathway that is independent of M1^14,16-18^. Moreover, neural inputs to the hand and forelimb representation in M1 mostly originate from PMd^19,20^, creating a cortico-cortical circuitry that has influences on motor network excitability^21-24^. Finally, where these cortico-cortical connections exist, the onset of movement-related activity occurs sooner in the premotor cortex than in M1^18,25^. As such, the premotor cortex may modulate motor output in two ways: first, through its dedicated CST projections to the spinal cord, and second, through its direct controlling connection to M1. Stimulating the premotor cortex could thus make the motor network more susceptible to cortical excitability changes. Only a few studies have examined the use of tDCS to modulate the premotor cortex, but most of them suggest that targeting the premotor cortex, and especially PMd, with tDCS could enhance motor performance and motor learning (for a review, see Buch & collaborators^26^). However, to our knowledge, there are no studies directly comparing the effects of tDCS over M1 versus the PMd on motor network excitability.

This lack of direct comparison is likely due to the poor spatial resolution of conventional tDCS montages. The distribution of the electrical current using a standard tDCS montage is relatively diffuse, leading to changes far beyond the stimulated regions^27^. Therefore, when M1 is targeted using conventional tDCS, at least a part of PMd^28^ is also likely stimulated. In contrast, high-definition tDCS (HD-tDCS) devices provide the ability to increase stimulation focality^29^, making it potentially easier to differentially target M1 versus PMd. Although there is still likely some overlap in the stimulation due to the anatomical proximity of these regions, the ring configuration of the HD-tDCS montage helps to concentrate the peak stimulation over the desired region.

In the current study, we focused on comparing (1) changes in cortical motor physiology and (2) changes in motor network connectivity and complexity across these three groups. Motor evoked potentials (MEP)^1^ were used to evaluate cortical motor excitability locally at the motor hotspot, while resting-state fMRI (rs-fMRI)^30^ was used to measure brain activity pre- and post-stimulation across the motor network. From the rs-fMRI data, we measured both functional connectivity of the motor network and multiscale entropy (MSE), which reflects the dynamics of neural networks across multiple time scales or temporal frequencies^31^ and the complexity of regional BOLD signals^32,33^. We hypothesized that applying HD-tDCS over the left premotor cortex would modulate motor network excitability more reliably than over the left motor hotspot in healthy individuals, and produce greater changes in motor connectivity and complexity, due to the additional motor pathways emerging from PMd. We also hypothesized that MSE should be more sensitive to tDCS-related changes than standard functional connectivity methods^33^, since MSE reflects the complexity of neural activity fluctuations associated with the probability of neural firing^33^ and has previously shown to be sensitive to excitability changes following anodal tDCS^32,34^. Lastly, as an exploratory analysis, we performed a post-hoc examination of the anatomical location of each individual’s stimulated area (as defined using single-pulse transcranial magnetic stimulation (TMS)) in relation to the assigned target brain area to see whether variability in the anatomical location of the stimulated area might relate to HD-tDCS responsiveness.

## RESULTS

### Demographics

Forty-six healthy participants took part in this double-blind, placebo-controlled study (Table 1). At inclusion, participants were randomly assigned to one of the following 3 groups: HD-tDCS applied over the left TMS-defined motor hotspot (Motor Hotspot, n=15), HD-tDCS applied over the left premotor cortex, which was defined as 2.5 cm anterior to the left motor hotspot (Premotor, n=15), or sham HD-tDCS (Sham, n=16, in which case, electrodes were evenly randomized over the left motor hotspot or the left premotor cortex). The primary experimenter was blind to the stimulation condition (sham versus real HD-tDCS) and only given the electrode location.

### Neurophysiological measurements

#### Baseline thresholds

We first checked whether participants in the Sham group showed any differences in the following baseline measurements based on electrode placement (i.e., over the left motor hotspot or the left premotor cortex): (1) Resting Motor Threshold (RMT), (2) stimulator intensity to induce an MEP of at least 0.75mV (TS0.75mV), or (3) pre-tDCS MEP amplitude. We used Student t-tests, and did not find any differences between individuals with motor hotspot versus premotor electrode positions across any parameters (RMT: t(14) = 0.68, *p* = 0.50; stimulator intensity to induce TS0.75mV: t(12) = 0.57, *p* = 0.58; pre-tDCS MEP amplitude: t(14) = 0.05, *p* = 0.96). We, therefore, combined the data to form a single Sham group, which was used in all the following analyses.

We then checked to see whether the initial cortical excitability (RMT, stimulator intensity to induce TS0.75mV) between the 3 groups was similar (Table 1). Using two separate one-way ANOVAs with “Group” (Motor Hotspot, Premotor, Sham) as a factor, we found that there were no significant differences between the 3 groups in the RMT (n = 45, Group effect: F(2) = 0.50, *p = 0*.61) or in the stimulator intensity used to reach TS0.75mV (n = 39, Group effect: F(2) = 0.49, *p* = 0.62).

#### Changes in MEP amplitude

##### Inter-individual variability in HD-tDCS response

Next, we examined differences in the reliability of tDCS to induce changes in cortical motor excitability across the 3 groups by assessing the number of individuals who responded to tDCS. We defined the “responders” as participants who showed an increase in MEP amplitude following HD-tDCS compared to baseline, as per previous research^12,35^ (see Methods section for more details). Using the Quantile Range Outliers method, we identified one outlier participant in the Motor Hotspot group (i.e., a change in MEP amplitude following HD-tDCS of more than 3 interquartile ranges, or more than 4.2mV; see Fig. 1). We removed this participant from further excitability analyses. In the Motor Hotspot group, 7 out of 14 (i.e., 50%) participants showed an increase in MEP amplitude, while in the Premotor group, 11 out of 15 participants (i.e., 73.3%) showed an increase in MEP amplitude (Fig. 1). In the Sham group, only 5 out of 16 participants (i.e., 31.25%) showed an increase in MEP amplitude following sham HD-tDCS, suggesting that some individuals will show some level of change in cortical excitability, even without stimulation. We also found considerable inter-individual variability in MEP changes following HD-tDCS across all the 3 groups (Fig. 1). The number of participants with increased MEP amplitudes post-HD-tDCS showed a trending difference across the 3 groups (χ^2^ (2) = 5.497, *p = 0*.06). Finally, as the boundary between responders and nonresponders using canonical definitions is very narrow^12,35^, we also explored different, more robust methods to classify responders/non-responders, which resulted in similar results. This additional analysis is summarized in Supplementary Data 1, Supplementary Table 2.

**Fig. 1:**
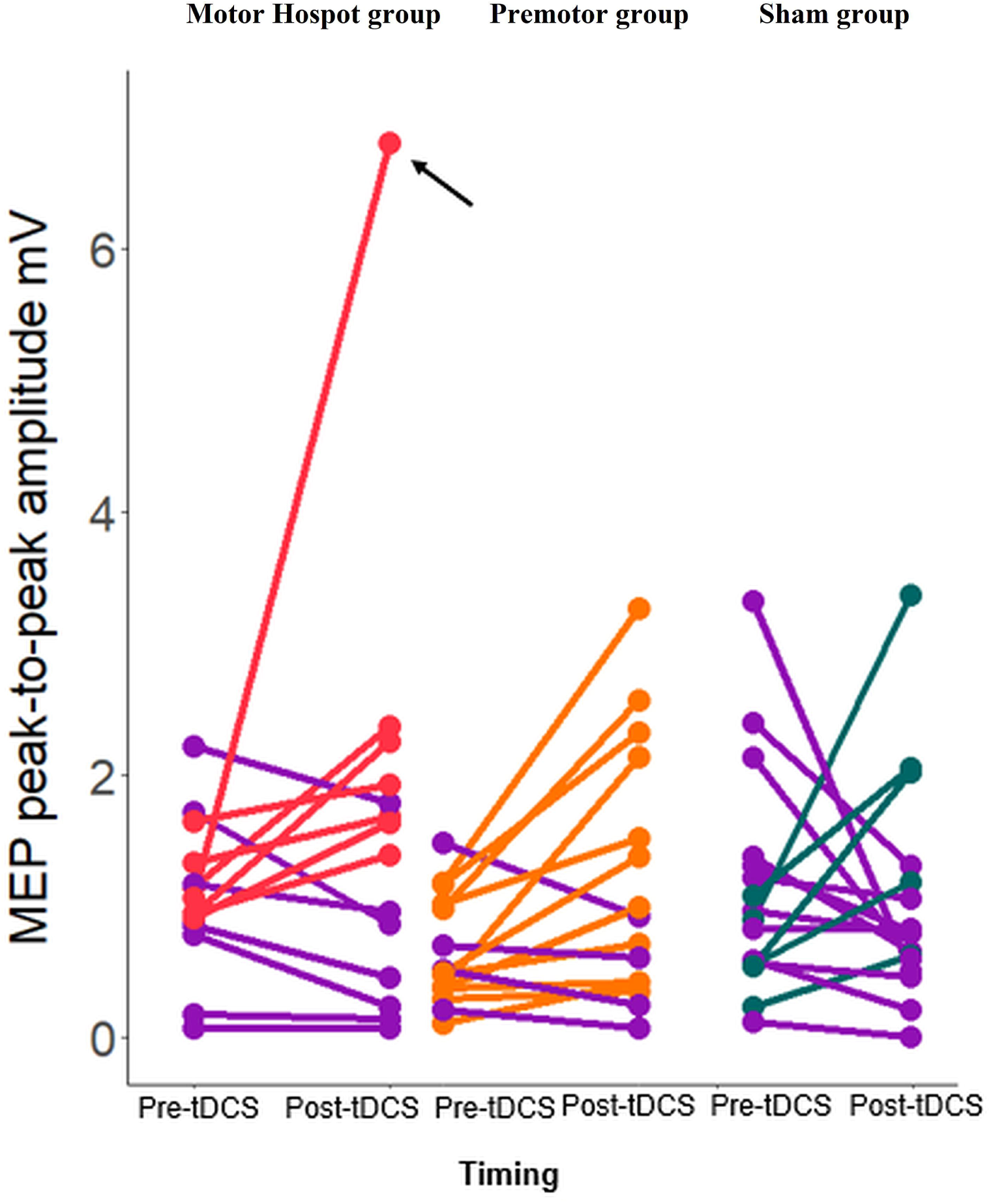
Individual changes in MEP amplitude over time. The dot pairs reflect MEP amplitude pre and post-tDCS for each participant in each of the 3 groups. Participants who showed a decrease in MEP between pre and post-tDCS are displayed in purple. *In the Motor Hotspot group, one participant presented a very strong effect post-tDCS. After running an outlier detection, this participant was removed from the rest of the analyses (indicated by the black arrow). *M1: primary motor area, MEP: motor evoked potential, mV: millivolts, PMd: dorsal premotor cortex, tDCS: transcranial direct current stimulation*.

##### Group level changes in MEP amplitude

Then, we examined whether HD-tDCS induced significant changes in cortical motor excitability across the 3 groups by comparing the MEP amplitudes before and after HD-tDCS (Table 1, Fig. 2).

**Fig. 2:**
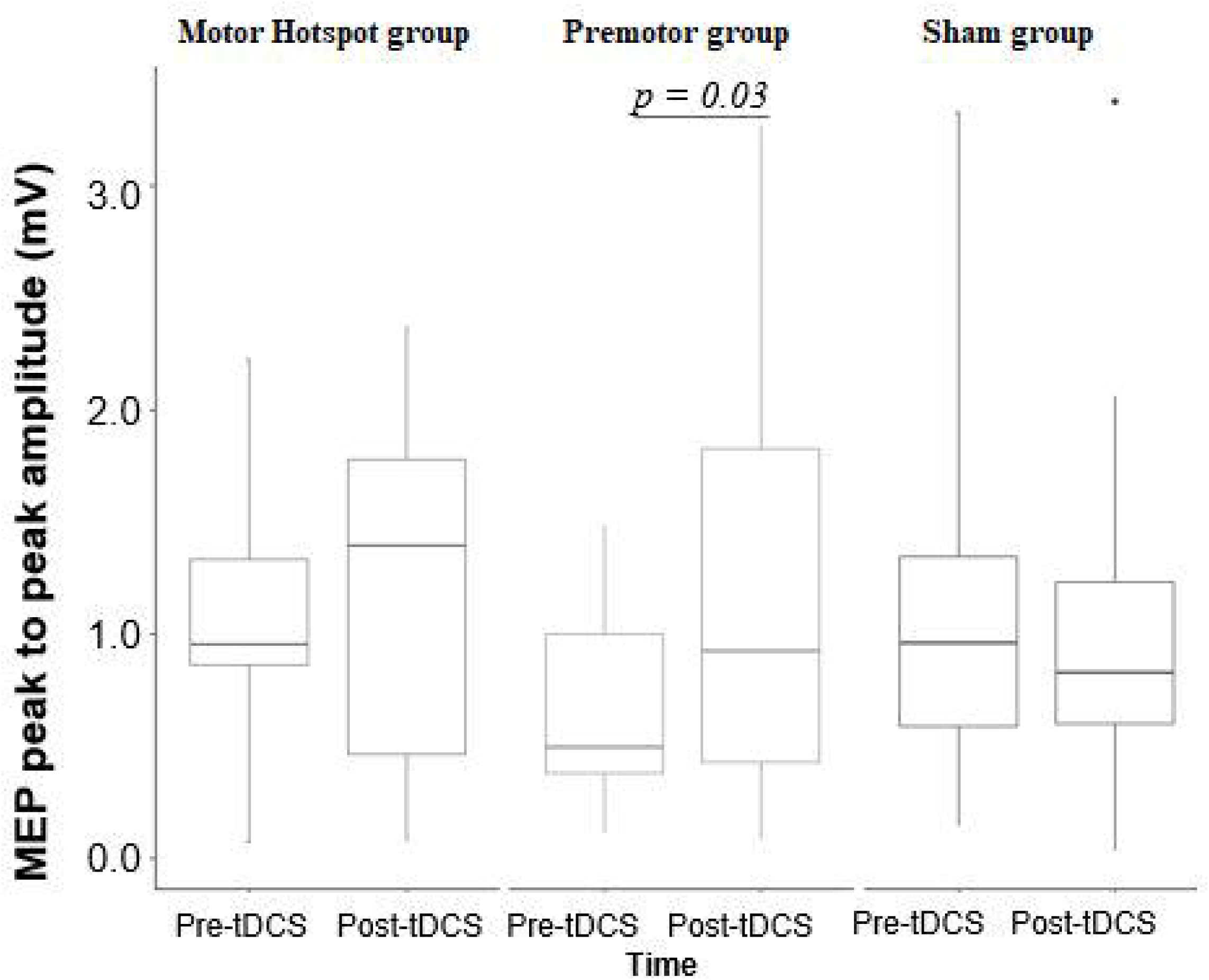
Changes in MEP amplitude for each group across time. Box-and-whisker plots of MEP amplitude per group over time (pre/post-tDCS). The center line represents the median value, the lower bound of the box represents the 25th percentile, the upper bound of the box the 75th percentile, and the whiskers represent 3 times the interquartile range. Any outliers are indicated by dots. Note that the only statistically significant difference is observed in the Premotor group reflecting an increase of MEP amplitude post-tDCS compared to pre-tDCS. The p-values refer to the statistically significant post-hoc contrasts. *M1: primary motor area, MEP: motor evoked potential, mV: millivolts; PMd: dorsal premotor cortex, tDCS: transcranial direct current stimulation*.

A two-way RM-ANOVA was performed on MEP amplitude, with “Time” (pre-HD-tDCS, post-HD-tDCS) and “Group” (Motor Hotspot, Premotor, Sham) as factors. There were no significant interaction or group effects (RM-ANOVA: n = 44, Interaction Group*Time F(2,41) = 1.88, *p =* 0.17, ƞ_p_^2^ = 0.31, Bayes Factor (BF) = 0.5 ± 1.29 %; Group: F(2,41) = 0.52, *p =* 0.59, ƞ_p_^2^ = 0.04, BF = 0.22 ± 0.54%; Time: F(1,41) = 1.83, *p =* 0.18, ƞ_p_^2^ = 0.0030, BF = 0.48 ± 1.13 %). Due to the large inter-individual variability, we then examined a priori comparisons of pre- versus post-HD-tDCS for each group, showing a significant increase in MEP amplitude only in the Premotor group (Premotor: post > pre-HD-tDCS t(41) = 2.56, p = 0.03, Cohen’s d = 0.73; Motor Hotspot: post > pre-HD-tDCS t(41) = 0.55, p = 0.58, Cohen’s d = 0.20; Sham post > pre-HD-tDCS t(41) = 0.44, p = 0. 66, Cohen’s d = 0.12).

### Resting-state fMRI

#### Resting-state functional connectivity

Next, we checked if HD-tDCS induced changes in rsfMRI motor network connectivity across the 3 groups by examining whether there were overall changes in the functional organization of the motor network. We defined four regions of interest (ROIs) as the left M1, right M1, left PMd and right PMd, using a meta-analysis from Hardwick and collaborators^36^ to define the location of the ROIs in the left hemisphere and then flipped them to the right hemisphere. We compared cross-correlation rankings for the six ROI-to-ROI pairs (1) left M1-right M1, 2) left PMd-right PMd, 3) left M1-left PMd, 4) left M1-right PMd, 5) left PMd-right M1, 6) right M1-right PMd) before versus after HD-tDCS.

The global organization of the motor network did not change after HD-tDCS in any of the groups, as demonstrated by a strong rank correlation within each group between pre and post-HD-tDCS measurements (Motor Hotspot: n = 15, r = 0.88, p = 0.03, Premotor: n = 15, r = 0.88, p = 0.03, Sham: n = 16, r = 0.94, p = 0.02). Moreover, there were also no significant differences in correlation ranks (Fisher’s Z test^37^) between groups (Motor Hotspot > Premotor: z = 0.0, p =1; Motor Hotspot > Sham: z = 0.78, p = 0.43, Premotor > Sham: z = 0.78, p = 0.43). This suggests that there was no global reorganization of the connectivity strengths in the motor network following HD-tDCS over either of the targets. We also found no significant changes post-HD-tDCS in the connectivity each ROI-to-ROI pair in any of the three groups (ANOVA, n = 46, thresholded at qFDR = 0.05; CONN toolbox). Additional exploratory analyses focusing on ROI-to-ROI comparisons and ICA network comparisons at a less stringent threshold (cluster-level corrected p<0.01) are reported in Supplementary Methods 1, Supplementary Data 2 & 3, Supplementary Fig. 1 & 2, Supplementary Table 1, and Supplementary Discussion.

#### Multi-scale entropy of resting-state fMRI

Then, we checked if HD-tDCS induced changes in motor network complexity by examining HD-tDCS-associated changes in MSE across the 3 groups (Fig. 3).

**Fig. 3:**
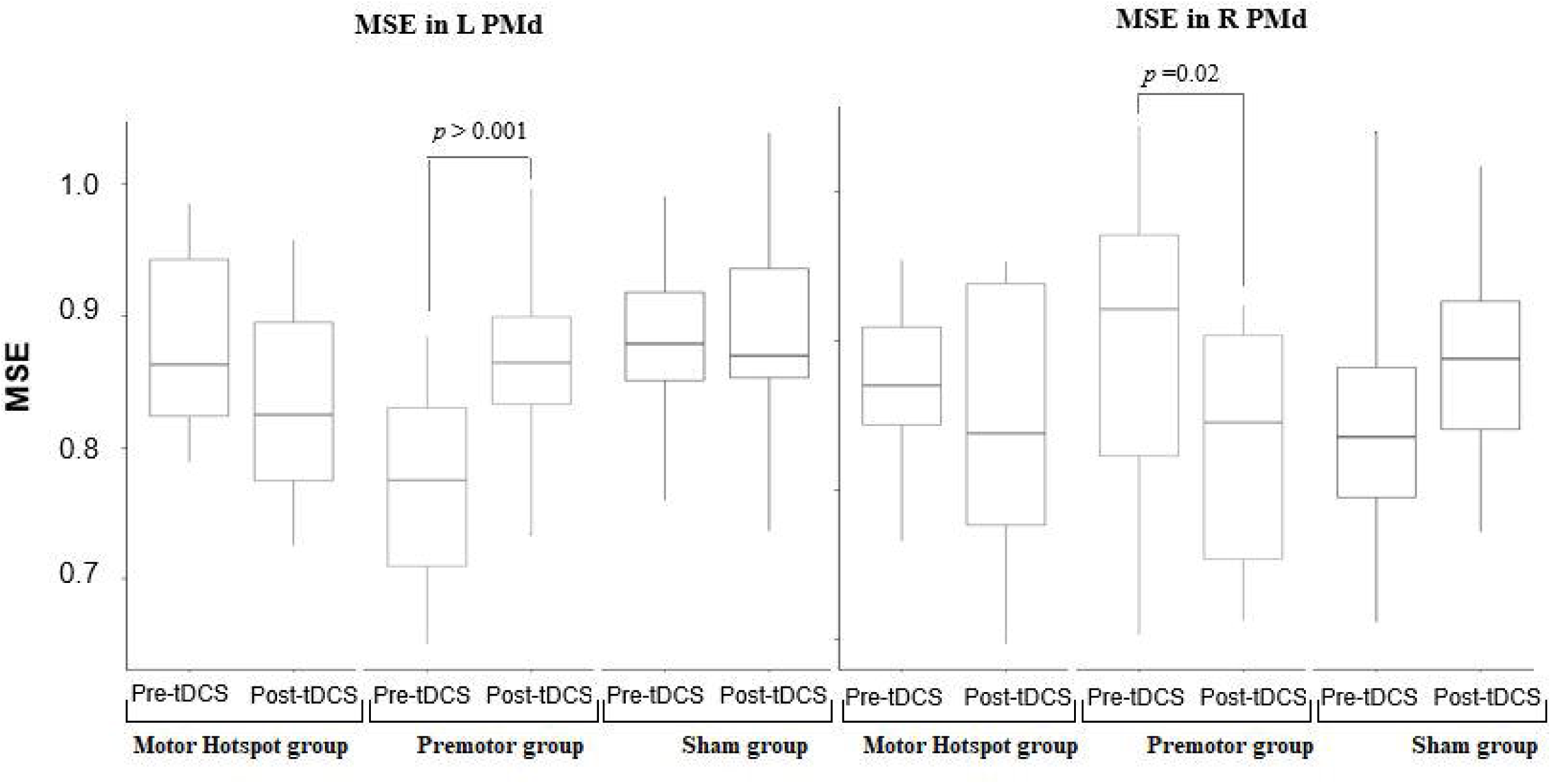
Changes in MSE across groups. Box-and-whisker plots of the MSE per group over time (pre/post-tDCS) in the left PMd ROI (in the left panel) and in the right PMd ROI (in the right panel). The center line represents the median value, the lower bound of the box represents the 25th percentile, the upper bound of the box the 75th percentile, and the whiskers represent 3 times the interquartile range. Any outliers are indicated by dots. The p-values refer to the statistically significant post-hoc contrasts. *L: left, M1: primary motor area, MSE: multiscale entropy, PMd: dorsal premotor cortex, R: right, tDCS: transcranial direct current stimulation*.

RM-ANOVAs with “Time” (pre-HD-tDCS, post-HD-tDCS) and “Group” (Motor Hotspot, Premotor, Sham) as factors were performed for each of the 4 regions of interest (left/right M1, left/right PMd) and demonstrated a significant interaction between group and time and significant main effect of group (Left PMd RM-ANOVA: n = 46, Interaction Group*Time F(2,43) = 9.06, p = 0.0005, ƞ_p_^2^= 0.14, BF = 498 ± 6.4 %; Group: F(2,43) = 7.5 p = 0.002, ƞ_p_^2^ 0.19, BF = 11.19 ± 0.61 %; Time: F(1,43) = 2.33, p = 0.13, ƞ_p_^2^ = 0.02, BF = 0.59 ± 1.91 %). Post-hoc analyses revealed a significant increase in MSE only in the left PMd following HD-tDCS over the premotor cortex (Premotor post > pre-HD-tDCS t(43) =4.09, p = 0.0002, Cohen’s d=1.31; Motor Hotspot post > pre-HD-tDCS t(43) =−1.94, p = 0.06, Cohen’s d=−0.61; Sham post > pre-HD-tDCS t(43) = 0.50, p = 0.61, Cohen’s d= 0.17).

In the right PMd, there was also a significant interaction and main effect of group (Right PMd RM-ANOVA: n = 46, Interaction Group*Time F(2,43) =3.89, p = 0.03, ƞ_p_^2^= 0.09, BF= 0.41 ± 2.85 %; Group: F(2,43) = 0.02, p = 0.9, ƞ_p_^2^ = 0.0003, BF= 0.14 ± 1.34 %; Time: F(1,43) =1.14, p = 0.29, ƞ_p_^2^= 0.009, BF= 0.45 ± 1.3 %). Post-hoc analyses revealed there was a decrease in right PMD MSE only in the Premotor group (Premotor post > pre-HD-tDCS t(43) =−2.43, p = 0.02, Cohen’s d = −0.78; Motor Hotspot post > pre-HD-tDCS t(43) =−0.71, p = 0.48, Cohen’s d=−0.28; Sham post > pre-HD-tDCS t(43) =1.48, p = 0.14, Cohen’s d= 0.60).

### Comparison of stimulation site location

Finally, given the high inter-individual variability of the results, we performed an additional post-hoc exploratory analysis to examine if the individual functional location of the left motor hotspot and the left premotor cortex matched the conventional/anatomical definitions for these regions (Fig. 4), and whether this could explain variability in HD-tDCS effects.

**Fig. 4:**
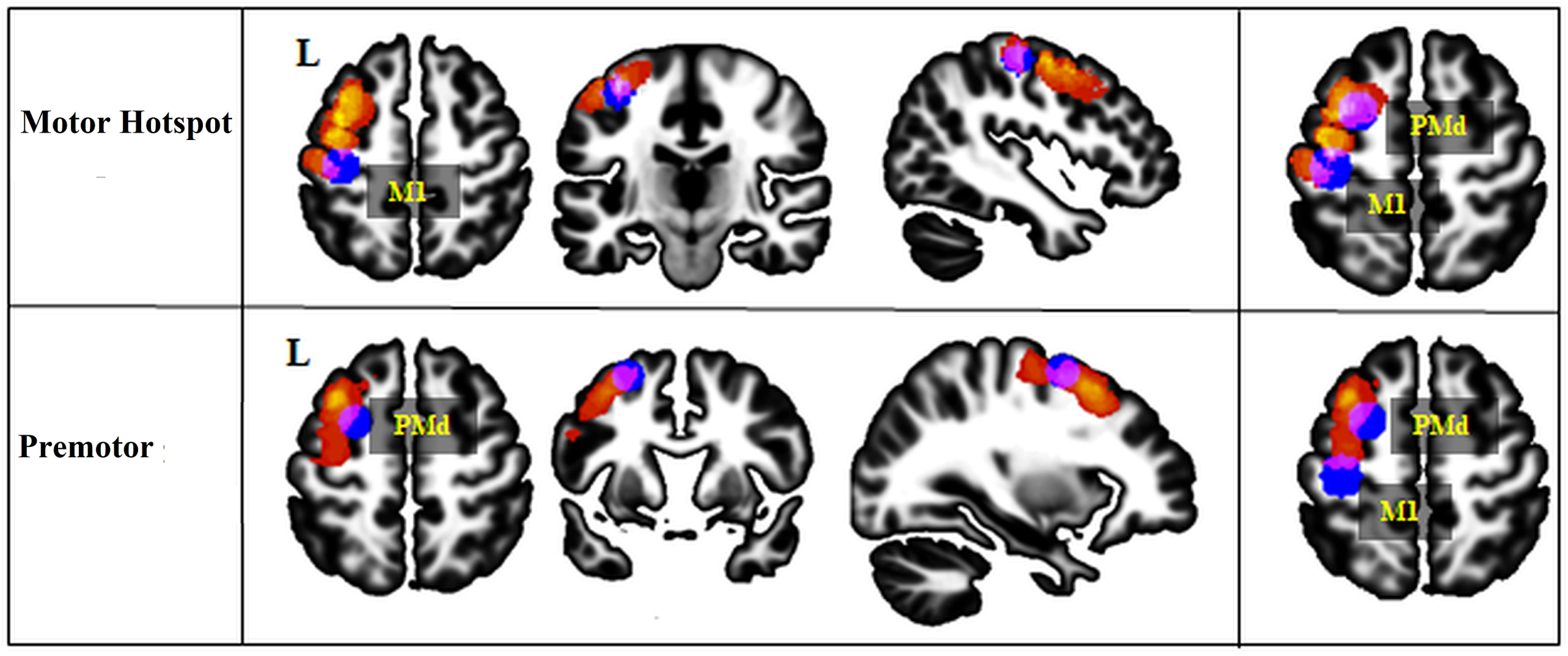
Location of the stimulation site. This figure displays the anatomical stimulation location sites in all the participants and shows an overlap of these sites with the standard left M1 or left PMd location for both groups. The anatomical stimulation sites are presented in a heat map, with individual subjects plotted in red and overlap between participants reflected in warmer colors (with yellow representing the surface shared by most of the participants). The standard M1 and PMd ROIs are displayed in blue. The overlap between the anatomical stimulation site location and the standard ROIs location is colored in purple. *M1: primary motor area, PMd: dorsal premotor cortex, L: left*.

In the Motor Hotspot group, the average minimal Hausdorff’s distance between individual’s cortex-projected stimulated region (i.e., the motor hotspot) and the standard M1 ROI was 3.79 ± 4.64mm, and with the standard PMd ROI, it was 3.4 6 ± 4.89mm. The motor hotspot site overlapped with the standard anatomical M1 ROI in 6 participants and with the standard anatomical PMd ROI in 6 participants. In the remaining 3 participants, the stimulation site was localized between the standard M1 and PMd ROIs.

In the Premotor group, the average Hausdorff’s distance between the anatomical location of the individual stimulation (i.e., premotor cortex stimulation) site and the standard PMd was 4.93 ± 4.80mm, and with the standard M1, it was 15.44 ± 4.78mm. The premotor cortex stimulation site overlapped with the standard PMd ROI in 5 participants and between the standard M1 and PMd ROIs in 2 participants. In the other 8 participants, the anatomical location of the stimulation site was localized slightly anterior to the standard PMd ROI, but still within the anatomical (atlas-based) boundaries of PMd.

Given the wide variability in the anatomical location of the stimulation site in the Motor Hotspot group, we reassigned the participants to groups based on the anatomical location of their stimulation site (see Supplementary Data 4, Supplementary Fig. 3). We confirmed our previous results, finding again a significant increase in MEP amplitude post-tDCS only in the newly defined anatomical PMd group (t(19)=2.28, p=0.03).

These results suggest that there is large inter-individual variability in the left motor hotspot and the premotor cortex anatomical localization using canonical methods employed in a majority of tDCS studies (e.g., TMS-defined motor hotspot ^6-10,12,38-44^ and PMd defined as 2.5 cm anterior to the motor hotspot ^45-48^). In particular, many individuals in the Motor Hotspot group had stimulation sites that were bordering or overlapping with the standard PMd ROIs, or somewhere between the standard M1 and PMd ROIs. This heterogeneity in the stimulation site could contribute to the variability of responsiveness to HD-tDCS.

We, therefore, compared the location of the stimulation sites between responders and non-responders. As shown in Fig. 5, more than half of the responders, including participants in both the Motor Hotspot and Premotor groups, received stimulation over the same area inside PMd anatomical boundaries. We found that 10 responders had an overlapping stimulation site within anatomical PMd region, regardless of group allocation (see Supplementary Fig. 4), while the anatomical location of the stimulation site in non-responders across groups was more widespread.

**Fig. 5:**
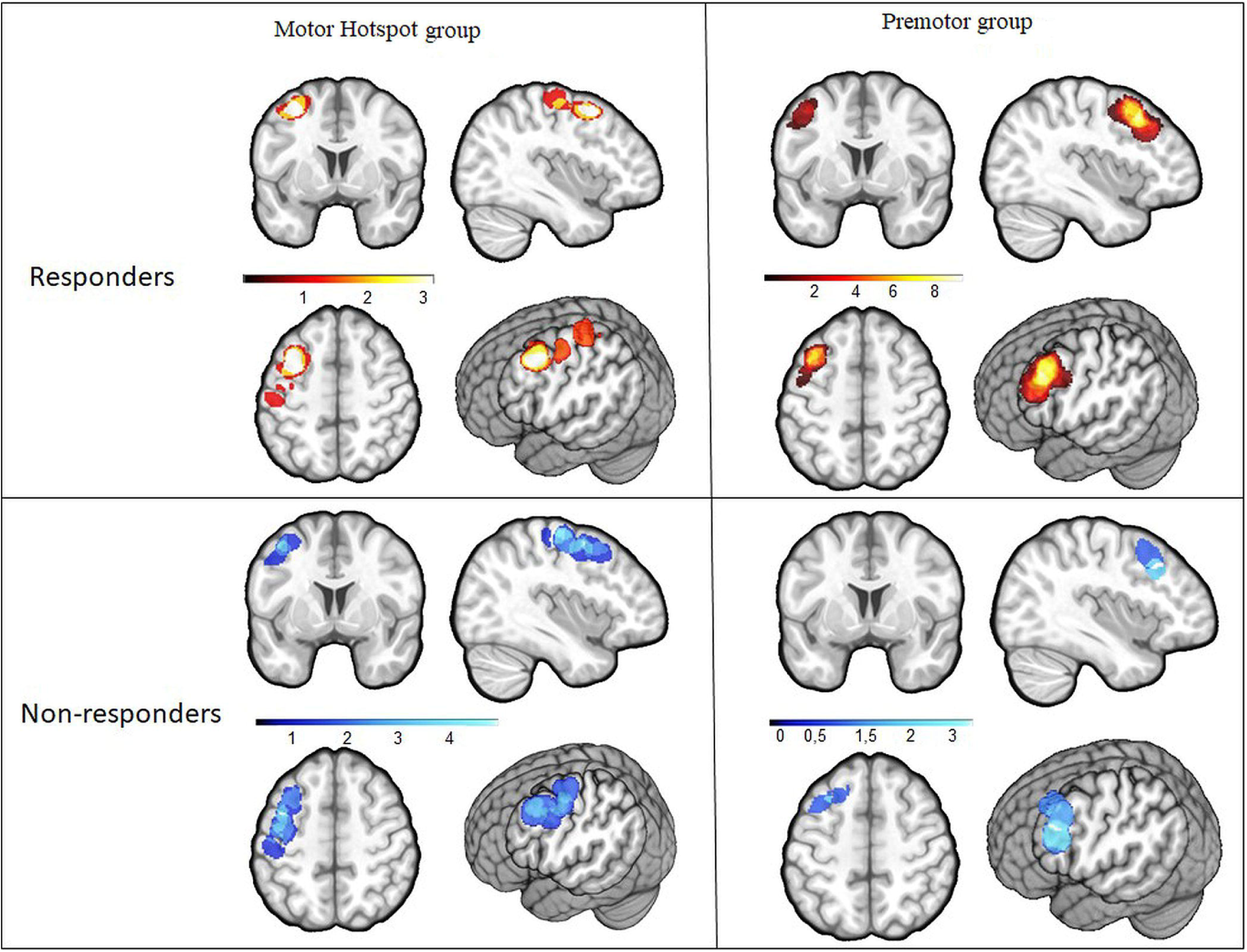
Heat map of the stimulation site between responders and non-responders in the Motor Hotspot and Premotor groups. This figure displays the location of the anatomical stimulation sites in the participants allocated to the Motor Hotspot group (n= 14) for the 7 responders (hot heat map) and the 7 non-responders (cool heat map), and to the Premotor group (n=15) for the 11 responders and the 4 non-responders. The majority of the responders in both Motor Hotspot and Premotor groups received stimulation over the same spot within the PMd anatomical boundaries (see also Supplementary Fig. 5).

## DISCUSSION

In this study, we performed a systematic, double-blind investigation using concurrent HD-tDCS and fMRI to examine changes in brain network connectivity and complexity after HD-tDCS stimulation of either the left motor hotspot (assumed to reflect the hand knob location in M1^3^) or the left premotor cortex. To our knowledge, this is the first known group study using concurrent HD-tDCS and fMRI to compare the left motor hotspot and left premotor cortex stimulation. We examined changes in cortical motor excitability using both TMS and fMRI after HD-tDCS. Finally, we explored inter-individual variability in the stimulation sites.

Overall, we observed that HD-tDCS applied over either the left motor hotspot or left premotor cortex produced highly variable changes across individuals. Due to the large inter-individual variability in response to tDCS, the magnitude of changes in cortical excitability was not significantly different between the 3 groups. However, despite this, we found that HD-tDCS over left premotor cortex produced more slightly reliable increases in cortical excitability than HD-tDCS over the left motor hotspot or sham (i.e., more participants showed increased MEP amplitude following HD-tDCS over left premotor cortex), and we found that MEP amplitude increased significantly after HD-tDCS only in the Premotor group. The slight increase of cortical excitability in the Premotor group was also accompanied by modulations in motor network complexity.

Finally, we showed that there was significant variability in the location of the anatomical stimulation sites compared to the expected stimulation sites based on canonical electrode placement – a factor which may have introduced variability into previous brain stimulation studies. In addition, the highest and most consistent number of responders was observed when stimulation occurred over left anatomical PMd. However, there was still large inter-individual variability in the stimulated anatomical location of responders, and some individuals in the non-responder group also were stimulated within PMd, suggesting that there is likely not a single anatomical target across all participants that would effectively increase excitability.

Altogether, these results suggest that HD-tDCS applied over premotor cortex may lead to small but more reliable changes in cortical excitability than over the motor hotspot. As previously observed, there was large variability in the MEP measurement even in the Sham group, suggesting that this measure is highly variable over time and across individuals. However, in line with our initial hypothesis, we observed a trend of a more reliable increase of MEP amplitude in the Premotor group, resulting in 75% responders compared to only 50% responders in the Motor Hotspot group (and 31% in the Sham group). Moreover, only the Premotor group showed a slightly significant increase post-tDCS. These results support previous findings^12^ that the motor hotspot may not be the best target to consistently enhance cortical excitability.

HD-tDCS over both the motor hotspot and premotor cortex led to no differences in motor network functional connectivity. Although previous work has suggested that tDCS may induce changes in functional connectivity of the motor network^49,50^, our analyses showed no significant changes. Our exploratory analyses, at a lower threshold, presented in Supplementary Data 2 and 3, Supplementary Fig. 1 and 2, and Supplementary Discussion, demonstrate a trend towards modifications in functional connectivity after HD-tDCS, but these modifications are only significant at lower statistical threshold. It is possible that the short duration of the HD-tDCS session (7 minutes) and low amplitude of current (1 mA) might not have been sufficient to induce reliable and significantly different changes in the functional connectivity of the motor network across individuals, which has been previously observed after 10 to 20 minutes of stimulation^49,50^.

On the other hand, we did find changes in multiscale entropy in left and right PMd following HD-tDCS over left premotor cortex. MSE, a method for measuring the complexity of finite length time series, quantifies the regularity of a time series by calculating the complexity of a signal at different time scales^51^. This measure reflects the dynamics of the neural networks^31^. Previous work has shown that increasing brain complexity is associated with brain adaptation^32,33^. Importantly, MSE is thought to measure fine-grained changes in brain function following tDCS^32,34,52^. Despite large variability in the baseline MSE measurements, HD-tDCS over premotor cortex, as hypothesized, led to significant and more reliable changes in MSE than HD-tDCS over the motor hotspot. Given that HD-tDCS over premotor cortex led to more reliable changes in cortical excitability, it is likely this was also captured in the MSE changes found in both hemispheres (i..e., an enhancement of the excitability in the stimulated hemisphere associated with a decrease of the excitability in the contralateral hemisphere^53,54^). In line with previous studies^33^, MSE and functional connectivity seemed to be complementary metrics: MSE may be more sensitive and able to capture local changes in the motor network whereas traditional rs-fMRI functional connectivity would capture more robust changes between distant brain areas.

In the present study, we also observed high inter-individual variability in HD-tDCS-induced changes in cortical excitability. This variability is similar to what has been recently reported, with up to 50% of participants showing no response to tDCS^11,44,55^. Here, we found a modest enhancement of the number of participants who responded to HD-tDCS when applied over premotor cortex. However, when we reallocated individuals based on their anatomical stimulation location, we still found a consistent increase in MEP amplitude only for the Premotor group (for those that received HD-tDCS over premotor cortex). We also consistently found the PMd to be the most-stimulated anatomical location in responders. This suggests that stimulating the premotor cortex to modulate the motor network might help reduce inter-individual variability compared to stimulating the motor hotspot, as long as it is actually the PMd that is being stimulated. Finally, our results suggest there is no single brain area that would ensure a positive response to tDCS across all participants. Although more than half of the participants who responded to tDCS received stimulation over an overlapping PMd spot, for the other half, the stimulation site was somewhere between the anatomical M1 and PMd. Moreover, in the participants who did not respond to tDCS, the stimulation site was all over the motor network, including in PMd, with no strong overlap across participants. Again, this suggests that more than the absolute location, differences in brain structure and function^56^ should be taken into account to determine the responsiveness of each individual to tDCS. Future studies should determine which stimulation parameters (neural target, nature of stimulation, duration, etc.) are the most effective to induce changes in motor network excitability and function at the individual level instead of following a one-size-fits-all method.

The current study also has several limitations. First, although the size of our groups (n=15-16 per group) is relatively large for studies looking at cortical excitability following brain stimulation, these groups are still relatively small, especially given the large variability within each group. Replicating these results in a larger sample would be useful. A second limitation is that the current Sham group included half motor hotspot and half the premotor cortex electrode positions, which showed no significant differences with one another, and which we, therefore, combined into a single Sham group. However, to provide proper control for each experimental group, a matched full Sham group for each experimental group should be used in the future. Third, we used a short 7-minute stimulation duration at 1 mA current, which may not have been enough to induce a strong effect, particularly in functional connectivity changes. However, as noted previously, this stimulation duration is in line with previous studies showing effects of 5-7 minutes of anodal tDCS on cortex excitability (for up to 30 min after the end of the stimulation)^57-59^. In addition, this study demonstrated that stimulating a brain region other than the motor hotspot (in this case, the premotor cortex) even for a short period can lead to an enhancement of cortical excitability and complexity, suggesting the duration was long enough to elicit some changes. However, future work should examine whole-brain network changes following longer durations or higher intensities of HD-tDCS. Forth, when turned on, the MRI-compatible HD-tDCS introduces noise into the BOLD signal with some undesirable, hard to remove, artifacts, that are even been observed in a cadaver^60^. Therefore, we were not able to analyze rs-fMRI data during brain stimulation—only immediately before and after. However, it would be very useful in future research to examine also BOLD signal changes during brain stimulation. Fifth, individual anatomy could have influenced the tDCS observed effect, as our anatomical findings suggest. Future studies that more carefully target stimulation based on individual anatomy, using each individual’s own anatomical brain scan with neuronavigation, would help address this issue. In addition, the present study does not include electrical field modeling, which is beyond the scope of the current investigation. However, in future studies, the impact of the anatomical properties of the stimulated area in the non-normalized brain of each individual could be used to explore the deeper impacts of the anatomical stimulation location on motor network excitability. Finally, we used a parallel study design, as opposed to a cross-over design. Each participant received HD-tDCS over only one neural target. Consequently, we cannot confirm that the participants who responded to HD-tDCS over premotor cortex would have not also responded to HD-tDCS over the motor hotspot, or vice-versa. Future studies involving a larger sample size and a cross-over design are needed to answer this specific question.

## Conclusion

To our knowledge, this is the first large study employing HD-tDCS with fMRI in a double-blind design with multiple stimulation sites. Using this design, we investigated the neurophysiological effects of tDCS on the motor network immediately after stimulation. We compared HD-tDCS over the motor hotspot versus the premotor cortex and showed that the premotor cortex, and especially PMd, may be a potential alternative neural target to modulate motor network excitability and neural complexity with less variability compared to HD-tDCS over the motor hotspot. These results also suggest that multiscale entropy may be a sensitive measure of changes in neuronal excitability following noninvasive brain stimulation within a single region and may be able to detect changes not observed with standard functional connectivity measures. Finally, we demonstrate that conventional methods for defining tDCS targets may be influenced by large inter-individual variability in the anatomy of the motor hotspot location. Future work should carefully consider and account for the anatomical, versus functional, stimulation locations. Altogether, these results open new considerations for improving the efficacy of tDCS on modulating the motor network.

## MATERIAL AND METHODS

### Population

The protocol was approved by the University of Southern California Institutional Review Board. All subjects gave written informed consent in accordance with the 1964 Declaration of Helsinki. The inclusion criteria were the following: (1) being a healthy volunteer aged 18 – 50 years, (2) being right-handed. Exclusion criteria were any contra-indications to MRI or brain stimulation.

### Study design

The study involved a single three-hour visit which included the following 3 sessions (see Fig. 6): (1) a pre-MRI TMS session, including the localization of the left motor hotspot and the left premotor cortex as well as the recording of neurophysiological measurements, (2) a concurrent HD-tDCS/MRI session, including the placement of the HD-tDCS cap, acquisition of structural MRI sequences, a pre-HD-tDCS rs-fMRI scan, the double-blinded HD-tDCS stimulation with concurrent acquisition of rs-fMRI scan, as well as a post-HD-tDCS rs-fMRI scan, and (3) a post-MRI TMS session to measure changes in neurophysiological measurements following HD-tDCS. To our knowledge, this is the first known randomized, double-blinded group study using a concurrent HD-tDCS and fMRI paradigm in humans. The safety of these concurrent designs was established in a pilot study using both phantoms and a single participant^61^. They also examined the tolerability of their specific double ring HD-tDCS montage in a behavioral session (without fMRI) in 30 participants.

**Fig. 6:**
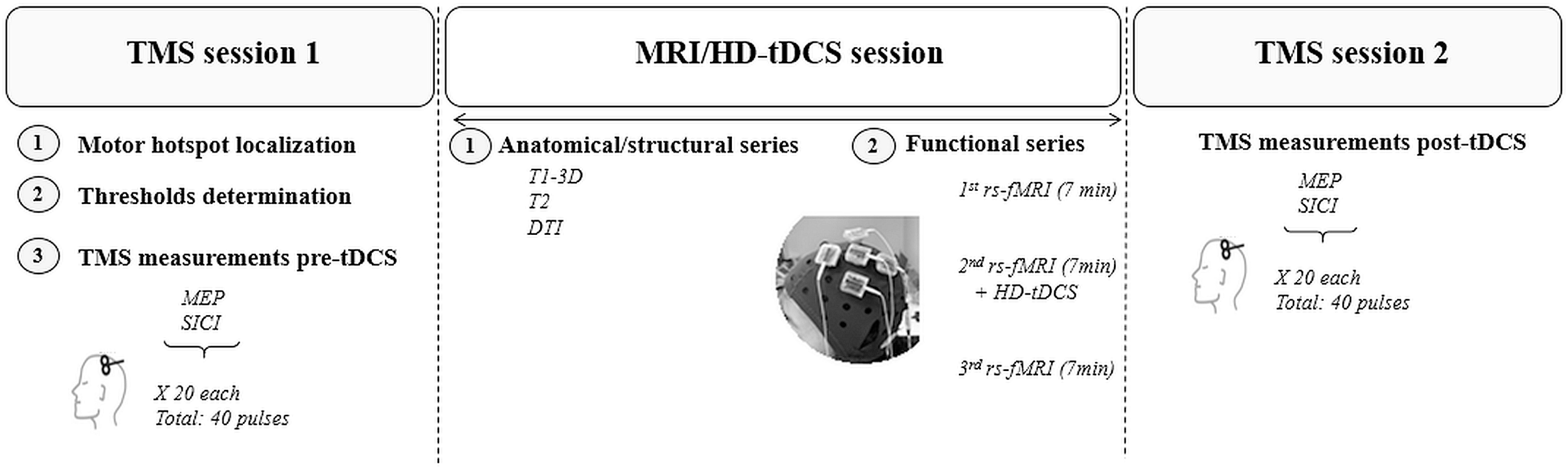
Study design. Each participant participated in 3 sessions within one day as follows: a first TMS session, then an MRI/HD-tDCS session, and finally a second TMS session. *HD-tDCS: high-definition transcranial direct current stimulation, MEP: Motor Evoked Potential, RMT: Resting Motor Threshold, rs-fMRI: resting-state functional magnetic resonance imaging*

#### Pre-MRI TMS session

Briefly, we used single- and paired-pulse transcranial magnetic stimulation (TMS) (Magstim 200² device with BiStim module; Magstim Inc., UK) with a figure-of-eight coil, along with the Brainsight neuronavigation system (Rogue Resolutions Ltd, UK) and surface electrodes to record muscle activity (electromyography, or EMG; Delsys Trigno wireless sensors, Delsys Inc, MA). To localize the motor hotspot, we recorded EMG at the right first dorsal interosseous (FDI). We then recorded pre-HD-tDCS cortical excitability, including each individual’s RMT and single-pulse motor-evoked potentials (MEPs) with a test stimulus at the intensity needed to generate an MEP of 0.75mV peak-to-peak amplitude. Notably, this test stimulus is in accordance with previous studies showing a range of test stimuli between 0.2mV and 4mV^62,63^. We also measured short-interval intracortical inhibition (SICI) using a paired-pulse TMS paradigm (see Brain Stimulation Intervention section below for details on the TMS paradigms). MEPs were recording using Signal 6.05 software (Cambridge Electronic Design Ltd., UK).

#### MRI session

Immediately following the first TMS session, participants were fitted with the HD-tDCS cap and began an MRI session including the acquisition of an anatomical image (T1w MPRAGE), and three 7-minute baseline rs-fMRI. Sham or HD-tDCS were applied during the second rs-fMRI. An additional experimenter, who was unblinded to the stimulation condition (real or sham), turned the tDCS stimulation device on and off. In this way, the primary experimenter and participant remained blinded to the stimulation condition. During the 3 rs-fMRI runs, participants were asked to look at a fixation cross and to not focus on any specific thoughts. The instructions were repeated at the beginning of each rs-fMRI run. Before the second rs-fMRI run, we informed the participants that the HD-tDCS stimulation was about to start, and we checked with them after the ramp-up to see if they were fine with the associated scalp sensations. Before the third rs-fMRI run, we let them know that the HD-tDCS had ended and that it would be the final rs-fMRI run.

#### Post-MRI TMS session

Immediately following the MRI session, the participants took part in a second TMS session to quantify the HD-tDCS-induced changes on cortical excitability. The cortical excitability measurements of this second TMS session started less than 10 minutes after the end of the third resting state fMRI scan.

We defined the “responders” subgroup as participants in the Motor Hotspot or Premotor group who showed an increase in MEP amplitude following HD-tDCS [(averaged MEP amplitude over the 20 trials post-tDCS) / (averaged MEP amplitude over the 20 trials pre-tDCS) > 1)]. The “non-responders” subgroup refers to participants in the Motor Hotspot or Premotor group who demonstrated a decrease or no change in MEP amplitude following HD-tDCS [(averaged MEP amplitude over the 20 trials post-tDCS) / (averaged MEP amplitude over the 20 trials pre-tDCS) ≤ 1)]. We also examined additional definitions of responders (see supplementary materials).

### Brain stimulation intervention

#### TMS paradigm

We defined the left motor hotspot using a canonical definition of the spot at which the maximal MEP peak-to-peak amplitude was recorded in the FDI muscle using a single TMS pulse at a given suprathreshold intensity (starting at 50% of the maximum stimulator output)^38-43^. At this location, we defined the RMT as the intensity of the stimulator that induced an MEP of at least 50 µV peak-to-peak amplitude in 5 out of 10 trials ^64^ and the TS0.75mV peak-to-peak amplitude in 5 out of 10 trials^62,63^.

Then, we acquired a MEP-SICI paradigm which consisted of 40 trials in a randomized order with a 7s ± 10% inter-trial duration. This paradigm included 20 single-pulse trials using the TS0.75mV (the peak-to-peak amplitude of these trials will be referred to as MEP amplitude), and 20 paired-pulse trials using a conditioning stimulus (CT) at 80% of the RMT 2.5ms before the TS0.75mV (HD-tDCS did not impact the amplitude of the SICI trials, for more details see Supplementary Data 5).

For 6 participants (2 in each group), we were unable to reach the intensity needed to elicit an MEP of 0.75mV, or even 0.5mV. In these cases, we used the RMT as the test stimulus to acquire MEP amplitude. For one additional participant in the Motor Hotspot group, we were only able to localize the motor hotspot but did not acquire any other neurophysiological measurements due to the participant’s reported discomfort. This participant was included only in the rs-fMRI analyses.

#### HD-tDCS experiment

We used the Soterix MRI-compatible HD-tDCS system (4×1 ring configuration) to deliver tDCS (1mA) for 7 minutes during an fMRI session (see Fig. 6). We used 5 small gel-filled electrodes (base diameter: 2.4 cm, gel-skin contact area of ∼4.5 cm^2^) positioned in adjacent electrode locations within the cap, and we placed the center electrode on the cap over the location of the neural target (as defined below). Each of the 4 cathodal ring electrodes delivered a −0.25mA current leading to a 1mA anodal stimulation over the center electrode.

#### Localization of the HD-tDCS targets

For the Motor Hotspot group, we positioned the center electrode over the left motor hotspot. For the Premotor group, we positioned the center electrode 2.5cm anteriorly to the motor hotspot, which is thought to represent PMd^65-69^. The use of this MRI-compatible device also gave us the unique opportunity to obtain an anatomical MRI with the exact position of the electrodes on the participant’s own head. We therefore also report the location of individual stimulation sites across participants in the Results section.

### MRI session acquisition and preprocessing

#### 2.4.1. MRI acquisition

MRI data were acquired on a 3T Prisma MRI scanner (Siemens, Germany) with a 32-channel head coil, using protocols from the Human Connectome Project (HCP) (http://protocols.humanconnectome.org/CCF/;https://www.humanconnectome.org/storage/app/media/documentation/lifespan-pilot/LSCMRR_3T_printout_2014.08.15.pdf). The MRI sequences acquired included the following: T1-weighted MPRAGE scan (208 sagittal slices, 0.8mm thick, TR = 2400ms, TE = 2.22ms, flip angle = 8°, voxel size 0.8 × 0.8 × 0.8mm); T2-weighted turbo spin-echo scan (208 sagittal slices, 0.8mm thick, TR = 3200ms, TE = 523ms, flip angle = 8°, voxel size 0.8 × 0.8 × 0.8mm); diffusion MRI scan (92 slices, 1.5 mm thick, TR = 3230ms, TE = 89.20ms, multi-band factor = 4, flip angle = 78°, voxel size 1.5 × 1.5 x1.5mm, with a gradient protocol with 7 scans at b=0 s/mm2, 47 at b=1500 s/mm2, 46 at b=3000 s/mm2, and a complete repetition with reversed phase encoding in the A-P direction); and three rs-fMRI sequences (7 minutes and 6 seconds covering 520 volumes of 72 slices per scan; TR = 800ms, TE = 37ms, flip angle = 52°, voxel size 2 × 2 × 2mm).

#### 2.4.2. Resting-state fMRI processing

We analyzed the rs-fMRI data using SPM12, the CONN toolbox (functional connectivity toolbox ^70^; http://www.nitrc.org/projects/conn) and the Complexity Toolbox (http://loft-lab.org/index-5.html). The preprocessing steps included slice-timing correction, motion realignment, noise correction using white matter, CSF and motion parameters as regressors, and band-pass filtering (0.01-0.1 Hz). We also performed co-registration between the functional scans and 3D-T1w MPRAGE scans of each participant. Finally, we normalized the functional scans to MNI space and smoothed them using a Gaussian filter of 6mm.

To measure functional changes in the motor network, we compared the functional connectivity between regions of interest (ROI-to-ROI analyses) and analyzed the neural complexity by computing the MSE of the BOLD time series in each ROI (defined below) before and after HD-tDCS. As mentioned in the results, we defined four ROIs as the left M1, right M1, left PMd and right PMd, using a meta-analysis from Hardwick and collaborators^36^ to define the location of the ROIs in the left hemisphere and then flipped them to the right hemisphere. The exact coordinates of the center of each ROI, which was defined as a sphere with a radius of 10mm, were: left M1 (x = −38, y = −24, z = 56), right M1 (x = 38, y = −24, z = 56), left PMd (x = −26, y = 2, z = 60), right PMd (x = 26, y = 2, z = 60).

##### ROI-to-ROI analyses

We extracted the cross-correlations of the 4 ROIs (left/right M1, left/right PMd) and ordered them for each group and each condition based on the correlation intensity. To compare the organization of the motor network following each stimulation session, we compared these orders using Spearman’s rank correlations coefficient.

We also used RM-ANOVAs (one RM-ANOVA for each ROI as a seed region) in the CONN toolbox to compare combinations of pairs between the 4 ROIs (the 6 functional connectivity pairs were: left M1-right M1, left PMd-right PMd, left M1-left PMd, left M1-right PMd, left PMd-right M1, right M1-right PMd) using “Time” (pre-HD-tDCS, post-HD-tDCS) and “Group” (Motor Hotspot, Premotor and Sham) as factors.

##### Multiscale-entropy

MSE was used to evaluate changes in neural complexity following HD-tDCS in each of the 4 ROIs (left M1, right M1, left PMd, right PMd). The MSE metric quantifies the regularity or complexity of biological signals across a range of coarse-grained timescales or temporal frequencies. Complex systems with 1/*f* power spectra exhibit constant entropy over various timescales (due to their fractal properties), whereas random noise shows a marked decrease in entropy at longer time scales (as random fluctuations are smoothed out)^33^. We used a pattern matching threshold (r) of 0.5 and a pattern length (m) of 2^33,71,72^. In total, we obtained 20 coarse-sampled scales. Based on the band-pass filter applied, we kept only 9 scales for the analyses (scales 12 to 20: frequency range [1/(TR*scale)]: 0.06-0.1 Hz).

### Statistical analyses

We used R (https://www.r-project.org/about.html) to perform all the statistical analyses, except for the analyses performed using the CONN toolbox (rs-fMRI functional connectivity). We used the CONN toolbox for ROI-to-ROI analysis, and the results are presented at a qFalse discovery rate = 0.05 (corrected for multiple comparisons).

We performed a repeated measures RM-ANOVAs to compare changes in MEP amplitude across groups, and separate RM-ANOVAS to compare changes in MSE for each of the 4 motor ROIs. We used factors of “Time” (pre-HD-tDCS, post-HD-tDCS) and “Group” (Motor Hotspot, Premotor and Sham) in all RM-ANOVAs. To evaluate HD-tDCS-induced changes within each group, we also performed a priori analyses of pre/post-HD-tDCS differences in MEP amplitude for each of the 3 groups using two-way paired t-tests. We report the test statistics and p-values, along with the effect size, using partial eta squared (ƞ_p_^2^) for the ANOVA main effects, and Cohen’s d for the t-tests. Any post-hoc tests are corrected for multiple comparisons using the Tukey HSD. In addition, given the large inter-individual variability in the data, we also reported the Bayes Factor (BF) for the group level changes over time in MEP amplitude and MSE analyses using the R package “BayesFactor”. The BF is used to report a ratio of the strength of the experimental effect compared to the null hypothesis and has recently been used with highly variable experimental data, including in brain stimulation studies^73,74^. In our data, we used the anovaBF to determine which model explained the best the variation observed in the results. We tested 3 models: impact of “Time”, impact of “Group” and impact of the interaction Group*Time. For each anovaBF, the model with the highest BF value was the be the best combination of models among these. The strength of the impact of this model on the data was determined by the size of the BF value. Briefly, a BF of 0-3 should be considered as weak evidence, a BF of 3-10 as substantial evidence, a BF of 10-30 as strong evidence and a BF >30 as very strong evidence^75^.

### Anatomical location of the stimulation site

We used the previously defined ROIs to define the standard M1 and PMd locations (see Fig. 7). The standard M1 ROI matched the location of the hand knob^3^, which is commonly thought to represent the anatomical location of the motor hand area. The standard PMd ROI was within the anatomical PMd boundaries (but due to the large extent of the anatomical PMd the functional ROI only covers a small portion of it).

**Fig. 7:**
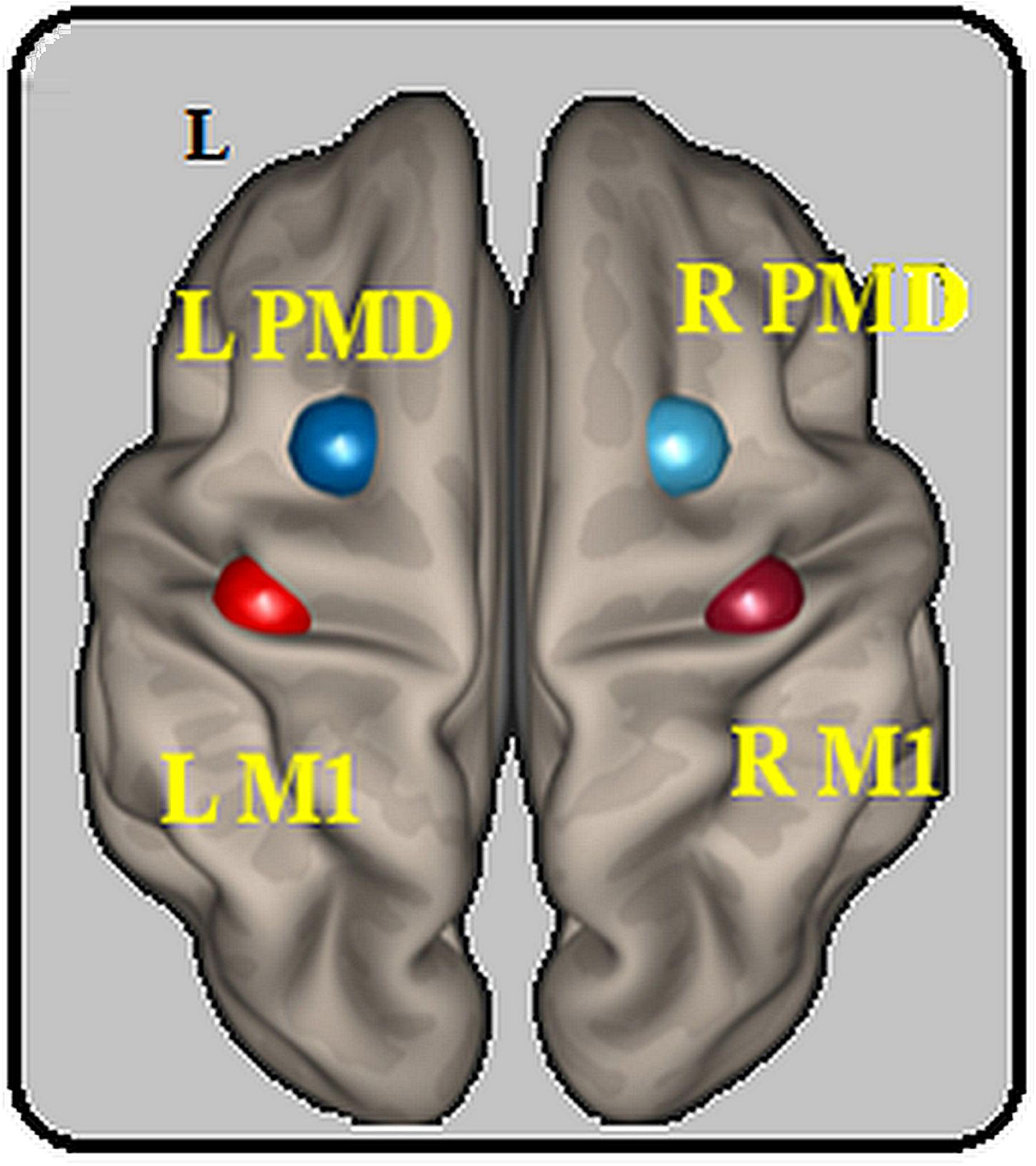
Location of the regions of interest. Location of the 4 ROIs based on the meta-analysis from Hardwick and collaborators^36^. Each ROI was defined as a sphere with a radius of 10mm around the reported peak coordinates for each region. These were: left M1 (x = −38, y = −24, z = 56), right M1 (x = 38, y = −24, z = 56), left PMd (x = −26, y = 2, z = 60), right PMd (x = 26, y = 2, z = 60). *L: left, M1: primary motor area, PMd: dorsal premotor cortex, R: right, ROI: region of interest*

In all the participants, the stimulation ROI was defined by a cortex-projection of the center electrode of the 4*1 ring. We projected the scalp coordinates of the center of this electrode on the MNI-normalized scans to the cortex. The distance of each voxel of the grey matter from “P” was calculated, enabling the creation of a spherical ROI centered at “P” (10mm diameter). This resulted in each participant with a cortex-projected stimulated region. We quantified the minimal distance of each individual’s cortex-projected stimulated region from the standard M1/PMd ROIs by computing the minimum Hausdorff’s distance (i.e., the minimum distance between the two ROI boundaries)^76^.

## Data availability

The unthresholded statistical maps used to generate the motor network (i.e., ICA analyses) are stored on the NEUROVAULT repository (https://neurovault.org/collections/UTZRKARR/). The raw data supporting these analyses will be made available following reasonable requests to the corresponding author. These data are not publicly available due to IRB restrictions.

## Fundings

This work was supported by our funding sources, including several National Institutes of Health grants that supported this work (NIH K01HD091283, R61MH110526 and R01MH111896) and the University of Southern California Provost’s Postdoctoral Scholar Research Grant.

## Acknowledgments

We would like to thank all the study participants, as well as Katherin Martin for her help during the MRI acquisition.

## Author contributions

S-L L and S.L. conceived and designed the study with input from D.J.J.W., K.J., and N.S.,

S.L., M.J. K.I., A.S. acquired the data,

S.L. M.J., K.J., and S.L.L. contributed to the data analyses and interpretation,

S.L. and S-L.L. wrote the paper with contributions from all the authors,

All the authors agreed with the submitted version or the paper.

## Competing interests

The authors declare no competing interests.

