## Supplementary material for "Differences in high-definition transcranial direct current stimulation over the motor hotspot versus the premotor cortex on motor network excitability"

**Supplementary Materials**

**Supplementary Data 1: Different ways to classify responders/non-responders to HD-tDCS**

One potential criticism of the current method for defining responders and non-responders as individuals with any MEP increase compared to baseline or decrease compared to baseline, respectively, is that even a very tiny magnitude of change could change the label of “responders” or “non-responders” if the mean is close to zero. Given the large variability of TMS responses within the same individual ^1^, the magnitude of change could be within that of the individual noise in the measurement. Therefore, we also tested several other measures of “response” that require a more stringent threshold to overcome any individual measurement noise (*Supplementary Table 1*).

A. *Using 10 % of the increase in MEP amplitude as a threshold.*

We first defined the “responders” as participants in the Motor Hotspot and Premotor groups who showed an increase in MEP amplitude higher than 10% of the pre-tDCS MEP amplitude [(averaged MEP’ amplitude over the 20 trials post-tDCS) > (110/100 * averaged MEP’ amplitude over the 20 trials pre-tDCS)]. The rationale for this decision was that the cut-off should exclude the spontaneous intra-individual fluctuation in MEP measure over time. The “non-responders” would be defined as follows: [(averaged MEP amplitude over the 20 trials post-tDCS) ≤ (110/100 * averaged MEP amplitude over the 20 trials pre-tDCS)].

Using this method, the response of the participants showing an MEP amplitude improvement is the same as the traditional cut-off used in the main paper. In the Motor Hotspot group, 7 out of 14 (50%) participants showed an increase in MEP amplitude, while in the Premotor group, 11 out of 15 participants (73.3%) showed an increase in MEP amplitude. In the Sham group, 5 out of 16 participants (31.25%) showed an increase in MEP amplitude following sham HD-tDCS (χ^2^ (2) = 5.497, *P =* 0.06).

B. *Using individual standard deviations (SD) to define the threshold*

We then defined the “responders” subgroup as participants in the Motor Hotspot and Premotor groups who showed an increase in MEP amplitude following HD-tDCS that was greater than their pre-tDCS MEP amplitude + one pre-tDCS SD (calculated at the individual level) [(individual averaged MEP amplitude over the 20 trials post-tDCS) > ((individual averaged MEP amplitude over the 20 trials pre-tDCS) + (individual pre-tDCS SD))]. The rationale for this decision was that the variability of the pre-tDCS assessment should reflect the spontaneous individual fluctuation in MEP amplitude. The “non-responders” would be defined as follows: [(individual averaged MEP amplitude over the 20 trials post-tDCS) ≤ ((individual averaged MEP amplitude over the 20 trials pre-tDCS) + (individual pre-tDCS SD pre-tDCS))].

Using this method, there were 4 individuals classified as responders in the Motor Hotspot group and 8 individuals classified as responders in the Premotor group. In the Sham group, 3 individuals also presented an increase of MEP amplitude. This method also shows more individuals responded to tDCS over The premotor cortex than the motor hotspot or Sham, although a chi-squared test did not show significantly different results (χ^2^ (2) = 4.37, *P =* 0.11).

C. *Using variability of the Sham group as a threshold.*

Then, we defined the “responders” subgroup as participants in in the Motor Hotspot and Premotor groups who showed an increase in MEP greater than the changes observed in the Sham group following HD-tDCS [(averaged MEP amplitude over the 20 trials post-tDCS) / ((averaged MEP amplitude over the 20 trials pre-tDCS) > (changes in MEP amplitude for each individual in the Sham group, averaged)]. The rationale for this decision was that changes in the Sham group should capture the measurement noise of the MEP measure since, in the absence of stimulation, changes in their MEP amplitude should just reflect normal variance. The “non-responders” would be defined as follows [(averaged MEP amplitude over the 20 trials post-tDCS) / ((averaged MEP amplitude over the 20 trials pre-tDCS) < (changes in MEP amplitude for each individual in the Sham group, averaged)].

In the Motor Hotspot group, 6 out of 14 participants showed an increase in MEP amplitude, while in the Premotor group, 11 out of 15 participants showed an increase in MEP amplitude. In the Sham group, 6 out of 16 participants showed an increase in MEP amplitude following sham HD-tDCS. However, these differences again were not significant (χ^2^ (2) = 4.53, *P =* 0.10).

Conclusion:

These analyses demonstrated that whatever the used cut-off was, there were more individuals classified as responders in the Premotor group than Motor Hotspot or Sham groups. Moreover, the 11 individuals classified as responders in the main paper, were still classified as responders with 2 out of 3 of the additional methods, suggesting a reliable response to the premotor cortex-tDCS.

**Supplementary Data 2: Whole brain ICA**

In order to evaluate tDCS-induced changes in functional connectivity of the whole brain, we performed an ICA analysis using the CONN toolbox with 20 ICs to compare the connectivity of five of the most common functional networks between the 3 groups over time. We compared the resting-state functional connectivity of the DMN, VN, CEN, SAL, and the motor network before versus after tDCS across the 3 groups (Group*Time interaction: F(2.43) = 3.21, *P*<0.01 cluster-size corrected). There were no significant changes over time in the DMN, the VN, the CEN or the SAL. In the motor network component, the Motor Hotspot group showed an increase of the resting-state functional connectivity post-tDCS compared with pre-tDCS in the right PMd. In addition, the Premotor group displayed an increase of the resting-state functional connectivity post-tDCS compared with pre-tDCS in the right parietal cortex. The Sham group showed no significant changes in the resting-state functional connectivity of the motor network over time. See also *Supplementary Figure 1, Supplementary Methods I, and Supplementary Discussion.*

**Supplementary Data 3: ROI-to-ROI Analyses**

In complement to the main analysis, we added an exploratory ROI-to-ROI analysis using a non-corrected threshold. This analysis demonstrated differences between pre and post-HD-tDCS only in the Motor Hotspot group (see *Supplementary Figure 2*). The comparison of before versus after tDCS showed a statistically significant enhancement of the resting-state functional connectivity between left PMd and right PMd (t(43) = 2.21). At that threshold, there were no significant differences in the rs-FC between the ROI pairs in the Premotor or Sham groups. For a discussion associated with these results, see *Supplementary Discussion.*

**Supplementary Data 4: Participant reallocation**

Using the anatomical location of the stimulation site, 6 participants of the Motor Hotspot group actually received stimulation over PMd. (No participants in the Premotor group actually received stimulation over M1.) Therefore, we re-analyzed the data from these individuals (however, we acquired pre/post-tDCS MEPs in only 5 of these participants). As displayed in Supplementary Figure 4, of these 5 participants, there were 3 participants with an increase of MEP amplitude over time (including the outlier), and 2 participants with a decrease of MEP amplitude over time. Using this reallocation, 20 participants were allocated to the new anatomical PMd group. Due to the imbalance of the groups after the re-allocation, we only performed Student paired t-tests to compare, in these anatomically allocated groups, MEP amplitude pre-tDCS (0.65 +/- 0.41 mV) to MEP amplitude post-tDCS (anatomical M1: 0.99 ± 0.44 mV vs 1.17 ± 0.85 mV, anatomical PMd: 0.79 ± 0.56 mV vs 1.5 ± 1.5 mV, Sham 1.15 ± 0.84 mV vs 1.05 ± 0.83 mV). We again found a significant increase of MEP amplitude post-tDCS only in the anatomical PMd group (anatomical PMd: t(19)=2.28, p=0.03, anatomical M1: t(8)=0.69, p=0.51, Sham: t(15)=-0.34, p=0.7).

**Supplementary Data 5: Effects of HD-tDCS on MEP amplitude in SICI trials**

To explore changes in intracortical inhibition, we compared MEP amplitude during the 0.75mV trials to MEP amplitude during the SICI trials (ratio of MEP amplitude divided by MEP amplitude in SICI trials) between the 3 groups before versus after tDCS.

As mentioned in the methods section, for 6 participants (2 in each group), we were not able to reach the S0.75mV threshold. Thus, we did not assess intracortical inhibition (SICI) in these 6 participants.

A two-way RM-ANOVA compared this ratio with “Time” (pre-tDCS, post-tDCS) and “Group” (Motor Hotspot, Premotor, Sham) as factors. There was no significant interaction effect and no significant main effect of time, but there was a nonsignificant trend for the main effect of Group (n= 39, interaction Group*Time: F(2,35) = 0.14*, P* = 0.87, ƞ_p_^2^ =0.002, Group: F(2,35) = 3.11, *P* = 0.06, ƞ_p_^2^ =0.11; Time: F(1,35) = 0.49, *P* = 0.49, ƞ_p_^2^ =0.004).

This analysis suggests that in the present study, neither HD-tDCS over the motor hotspot nor over the premotor cortex significantly affected intracortical inhibition, although there was a non-significant trend for the Sham group to show a smaller SICI overall than the two other groups (no changes over time).

**Supplementary Figure 1**

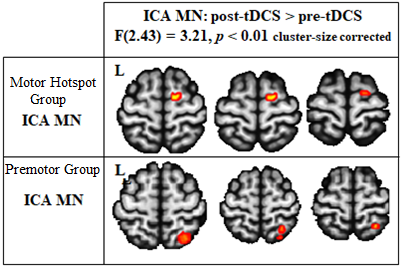
 **Supplementary Figure 1: Comparison of the ICA motor network over time**. The results of the whole brain ICA motor network comparison (P < 0.01, cluster-size corrected) are the following: the Motor Hotspot group showed an increase of the resting-state functional connectivity post-tDCS compared with pre-tDCS in the right PMd and the Premotor group in the right parietal cortex.

*ICA: independent component analysis, MN: motor network, rs-FC: resting-state functional connectivity, tDCS: transcranial direct current stimulation.*

**Supplementary Figure 2**

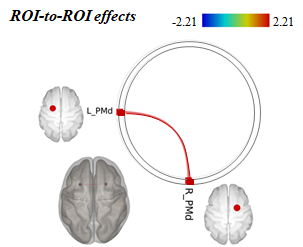

**Supplementary Figure 2: Comparison of the ROI-to-ROI resting-state functional connectivity.** The comparison of the ROI-to-ROI correlation between the 4 ROIs (6 ROI pairs) across the 3 groups (t(43) = 2.21), leads only to a slightly statistically significant enhancement of the resting-state functional connectivity in the Motor Hotspot group between left PMd and right PMd post-tDCS.

*L: left, PMd: dorsal premotor cortex, R: right, ROI: Region of interest.*

**Supplementary Figure 3**

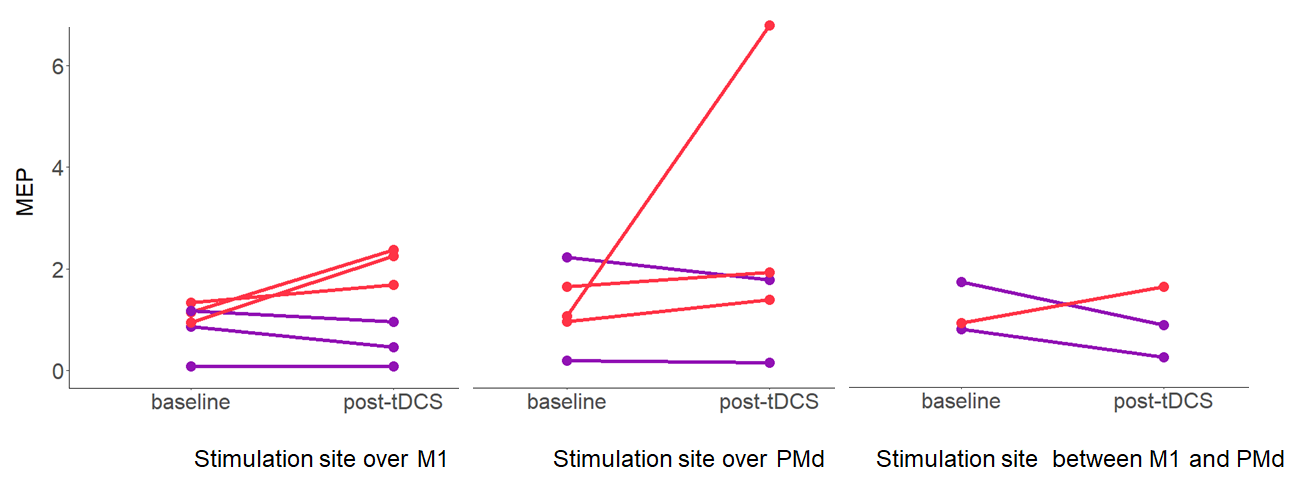
 **Supplementary Figure 3: Individual changes in the Motor Hotspot group’s MEP amplitude based on anatomical stimulation site.** The dot pairs reflect MEP amplitude pre and post-tDCS for the participants assigned to the Motor Hotspot group with an anatomical stimulation site over M1 (left panel), over PMd (middle panel) and between M1 and PMd (right panel). *MEP: motor evoked potential, PMd: dorsal premotor cortex, tDCS: transcranial direct current stimulation.*

**Supplementary Figure 4**

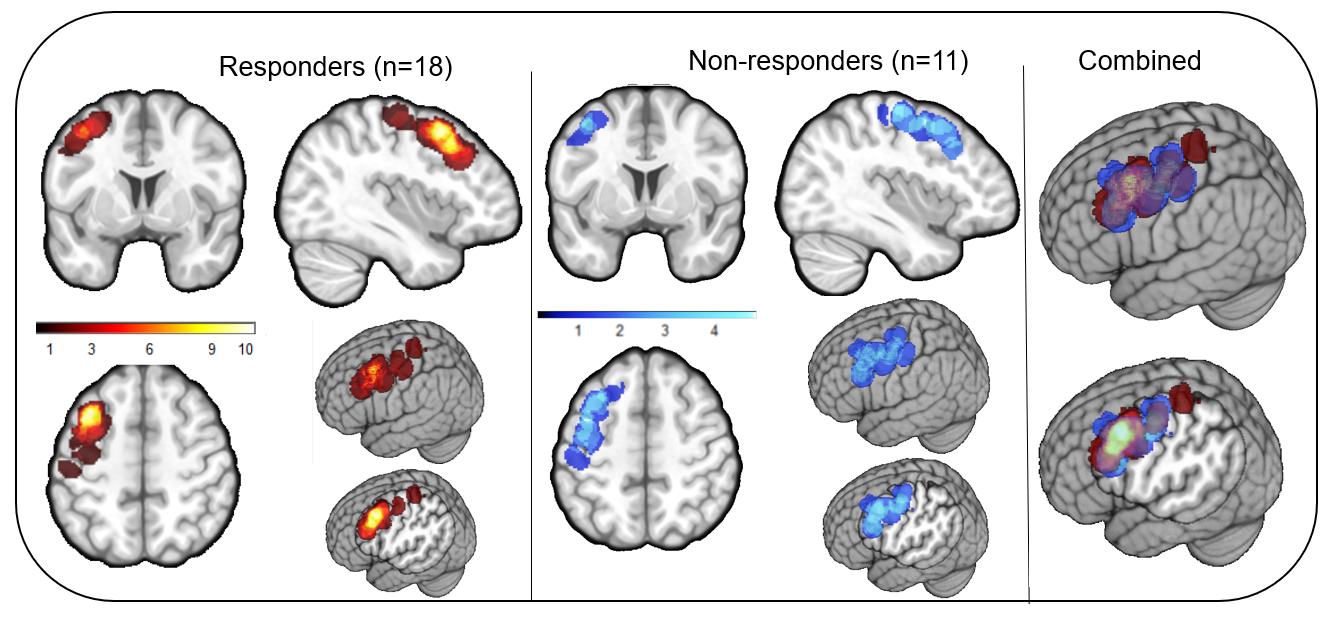
 **Supplementary Figure 4: Heat map of the stimulation site between responders and non-responders.** This figure displays the localization of the actual stimulation sites in the 18 responders (hot heat map) and the 11 non-responders (cool heat map) across both groups. Up to 10 participants shared the same stimulation site in the responder subgroup, whereas no stimulation sites were shared by more than 4 participants in the non-responders’ subgroup.

***Supplementary Table 1***

| ***Subject*** | ***Group*** | ***MEP pre-tDCS (mV)*** | ***SD pre-tDCS*** | ***MEP post-tDCS (mV)*** | ***Ratio MEP (post/pre)*** | ***Calcul threshold mean +10%*** | ***Responders Cut-off (ratio MEP>1 )*** | ***Responders Cut-off (post>pre+10%)*** | ***Responders Cut-off (averaged ratio changes in group 3)*** | ***Responders Cut-off (post> pre+ SD)*** |
| --- | --- | --- | --- | --- | --- | --- | --- | --- | --- | --- |
| *s01* | *Motor Hotspot* | *1.14* | *1.10* | *2.37* | *2.08* | *1.24* | *1* | *1* | *1* | *1* |
| *s02* | *Motor Hotspot* | *--* | *--* | *--* | *--* | *--* | *--* | *--* | *--* | *--* |
| *s03* | *Motor Hotspot* | *1.72* | *0.44* | *0.87* | *0.51* | *1.82* | *0* | *0* | *0* | *0* |
| *s04* | *Motor Hotspot* | *1.33* | *0.38* | *1.68* | *1.26* | *1.43* | *1* | *1* | *1* | *0* |
| *s05* | *Motor Hotspot* | *2.22* | *1.14* | *1.78* | *0.80* | *2.32* | *0* | *0* | *0* | *0* |
| *s06* | *Motor Hotspot* | *1.17* | *0.62* | *0.96* | *0.82* | *1.27* | *0* | *0* | *0* | *0* |
| *s07* | *Motor Hotspot* | *0.86* | *0.59* | *0.46* | *0.53* | *0.96* | *0* | *0* | *0* | *0* |
| *s08* | *Motor Hotspot* | *0.94* | *0.41* | *2.25* | *2.39* | *1.04* | *1* | *1* | *1* | *1* |
| *s09* | *Motor Hotspot* | *0.79* | *0.94* | *0.24* | *0.30* | *0.89* | *0* | *0* | *0* | *0* |
| *s10* | *Motor Hotspot* | *0.07* | *0.00* | *0.07* | *1.00* | *0.17* | *0* | *0* | *0* | *0* |
| *s11** | *Motor Hotspot* | *1.06* | *0.62* | *6.8* | *6.42* | *1.16* | *1* | *1* | *1* | *1* |
| *s12* | *Motor Hotspot* | *0.18* | *0.00* | *0.15* | *0.83* | *0.28* | *0* | *0* | *0* | *0* |
| *s13* | *Motor Hotspot* | *1.64* | *0.93* | *1.92* | *1.17* | *1.74* | *1* | *1* | *1* | *0* |
| *s14* | *Motor Hotspot* | *0.91* | *0.45* | *1.63* | *1.79* | *1.01* | *1* | *1* | *1* | *1* |
| *s15* | *Motor Hotspot* | *0.95* | *0.50* | *1.39* | *1.46* | *1.05* | *1* | *1* | *1* | *0* |
| *Mean* | | *1.07 (*1.07)* | *0.31* | *1.21(*1.61)* | *1.53* | *1.17* | *R=7* | *R=7* | *R=6* | *R=4* |
| *SD* | | *0.58 (*0.56)* | *0.14* | *0.80(*1.68)* | *1.53* | *0.56* | *NR =7* | *NR =7* | *NR =8* | *NR =10* |
| *s16* | *Premotor* | *1.18* | *0.38* | *3.26* | *2.76* | *1.28* | *1* | *1* | *1* | *1* |
| *s17* | *Premotor* | *0.48* | *0.73* | *0.71* | *1.48* | *0.58* | *1* | *1* | *1* | *0* |
| *s18* | *Premotor* | *0.38* | *0.51* | *0.49* | *1.29* | *0.48* | *1* | *1* | *1* | *0* |
| *s19* | *Premotor* | *1.48* | *0.33* | *0.92* | *0.62* | *1.58* | *0* | *0* | *0* | *0* |
| *s20* | *Premotor* | *0.98* | *0.88* | *2.56* | *2.61* | *1.08* | *1* | *1* | *1* | *1* |
| *s21* | *Premotor* | *0.38* | *0.18* | *0.99* | *2.61* | *0.48* | *1* | *1* | *1* | *1* |
| *s22* | *Premotor* | *1.17* | *0.00* | *2.32* | *1.98* | *1.27* | *1* | *1* | *1* | *1* |
| *s23* | *Premotor* | *0.3* | *0.34* | *0.41* | *1.37* | *0.40* | *1* | *1* | *1* | *0* |
| *s24* | *Premotor* | *0.11* | *0.00* | *0.43* | *3.91* | *0.21* | *1* | *1* | *1* | *1* |
| *s25* | *Premotor* | *1.02* | *0.38* | *1.52* | *1.49* | *1.12* | *1* | *1* | *1* | *1* |
| *s26* | *Premotor* | *0.21* | *0.58* | *0.08* | *0.38* | *0.31* | *0* | *0* | *0* | *0* |
| *s27* | *Premotor* | *0.39* | *0.41* | *2.13* | *5.46* | *0.49* | *1* | *1* | *1* | *1* |
| *s28* | *Premotor* | *0.52* | *0.18* | *0.25* | *0.48* | *0.62* | *0* | *0* | *0* | *0* |
| *s29* | *Premotor* | *0.7* | *1.35* | *0.61* | *0.87* | *0.80* | *0* | *0* | *0* | *0* |
| *s30* | *Premotor* | *0.49* | *0.00* | *1.38* | *2.82* | *0.59* | *1* | *1* | *1* | *1* |
| *Mean* | | *0.68* | *0.70* | *1.20* | *2.01* | *0.75* | *R=11* | *R=11* | *R=11* | *R=8* |
| *SD* | | *0.41* | *0.59* | *0.96* | *1.39* | *0.41* | *NR=4* | *NR=4* | *NR=4* | *NR=7* |
| *s31* | *Sham* | *1.39* | *0.45* | *0.68* | *0.49* | *1.49* | *0* | *0* | *0* | *0* |
| *s32* | *Sham* | *0.26* | *0.89* | *0.65* | *2.5* | *0.36* | *1* | *1* | *1* | *0* |
| *s33* | *Sham* | *2.14* | *0.41* | *0.62* | *0.29* | *2.24* | *0* | *0* | *0* | *0* |
| *s34* | *Sham* | *0.99* | *0.21* | *0.82* | *0.83* | *1.09* | *0* | *0* | *0* | *0* |
| *s35* | *Sham* | *0.61* | *0.40* | *0.23* | *0.38* | *0.71* | *0* | *0* | *0* | *0* |
| *s36* | *Sham* | *2.41* | *0.46* | *1.33* | *0.55* | *2.51* | *0* | *0* | *0* | *0* |
| *s37* | *Sham* | *0.92* | *0.70* | *3.37* | *3.66* | *1.02* | *1* | *1* | *1* | *1* |
| *s38* | *Sham* | *0.58* | *0.00* | *1.2* | *2.07* | *0.68* | *1* | *1* | *1* | *1* |
| *s39* | *Sham* | *0.85* | *0.64* | *0.85* | *1* | *0.95* | *0* | *0* | *0* | *0* |
| *s40* | *Sham* | *0.59* | *1.17* | *0.49* | *0.83* | *0.69* | *0* | *0* | *0* | *0* |
| *s41* | *Sham* | *1.33* | *0.27* | *0.83* | *0.62* | *1.43* | *0* | *0* | *0* | *0* |
| *s42* | *Sham* | *0.14* | *0.25* | *0.03* | *0.21* | *0.24* | *0* | *0* | *0* | *0* |
| *s43* | *Sham* | *1.24* | *1.08* | *0.87* | *0.70* | *1.34* | *0* | *0* | *1* | *0* |
| *s44* | *Sham* | *3.32* | *0.53* | *0.16* | *0.05* | *3.42* | *0* | *0* | *1* | *0* |
| *s45* | *Sham* | *0.56* | *2.03* | *3.63* | *6.48* | *0.66* | *1* | *1* | *0* | *1* |
| *s46* | *Sham* | *1.1* | *2.06* | *1.87* | *1.70* | *1.2* | *1* | *1* | *1* | *0* |
| *Mean* |  | *1.15* | *1.05* | *1.10* | *1.25* | *1.25* | *R=5* | *R=5* | *R=6* | *R=3* |
| *SD* | | *0.84* | *0.83* | *1.04* | *1.09* | *0.84* | *NR=11* | *NR=11* | *NR=10* | *NR=13* |

***Supplementary Table 1: Individual data and threshold for responders versus non-responders***

**Supplementary Table 2**
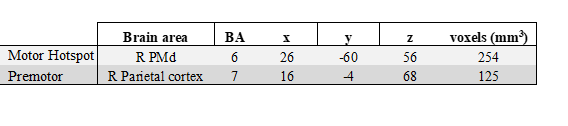

**Supplementary Table 2: Results of the ICA motor network comparison over time in the Motor Hotspot and Premotor groups (P<0.01, cluster-size corrected).**

**Supplementary Discussion:**

The functional connectivity modifications observed in the exploratory analyses of the functional connectivity (ROI-to-ROI and ICA analyses using a non-stringent threshold) were only observed at less stringent thresholds. First, we found that functional connectivity within the motor network was increased in the right hemisphere following tDCS of the left hemisphere (Motor Hotspot or Premotor groups). Specifically, in the ICA analysis, when looking at the motor network component, HD-tDCS over the motor hotspot led to a functional connectivity increase in the right PMd whereas HD-tDCS over The premotor cortex led to a functional connectivity increase in the right parietal cortex. In the ROI-to-ROI analysis, however, only the Motor Hotspot group showed an increase in left-right PMd connectivity. This M1 influence on interhemispheric PMd connectivity, but not on M1 connectivity, is confusing. However, one potential explanation for this is the anatomical localization of the actual stimulation site in the Motor Hotspot group, where many of the participants actually received stimulation closer to PMd. More research would be needed to better understand this finding. In the Premotor group, only the ICA analysis showed an increase in right parietal cortex. As the parietal cortex was not an ROI used in the ROI-to-ROI analyses, it remains to be seen whether this would have been a consistent finding. However, PMd-parietal cortex functional connectivity is a well-known pathway involved in the fronto-parietal reaching control network ^2^, so there is some basis for this modulation. Overall, however, we found that the functional connectivity results were very weak and difficult to understand. As noted in the main discussion, the duration of the stimulation may have been too short or weak to induce significant modulations in the connectivity. To better understand network changes following the motor hotspot versus The premotor cortex stimulation, future studies should provide longer stimulation to look at network connectivity changes.

**Supplementary Methods 1: Resting-state functional connectivity - Independent component analysis (ICA)**

As an exploratory analysis, we extracted 20 independent components (ICs) per participant (following Calhoun and colleagues’ ICA methodology ^3^) and then ran a fastICA algorithm allowing for a group-level independent component definition that accepts only one component per subject in each cluster. Across the 20 ICs, we identified five networks of interest ^4^^,^^5^: (1) the Default Mode Network (DMN) ^6^ including the posterior cingulate cortex, medial prefrontal cortex, lateral parietal cortex, and parahippocampal gyrus; (2) the central executive network (CEN) ^7^ including the dorsolateral prefrontal cortex and the posterior parietal cortex; (3) the motor network ^8^ including the primary and higher-order motor, sensory and parietal areas; (4) the salience network (SAL) ^7^ including the anterior insula and dorsal anterior cingulate; and (5) the visual network (VN) ^4^ including much of the occipital cortex. Using an ANOVA in the SPM CONN toolbox, we compared the spatial components of each of the aforementioned 5 networks using “Time” (pre-tDCS, post-tDCS) and “Group” (Motor Hotspot, Premotor, Sham) as factors. As this was an exploratory analysis, the results of this analysis are reported using a more liberal threshold (P<0.01 cluster-size corrected). Results are reported in *Supplementary Data 2*.

1	Schambra, H. M., Ogden, R. T., Martinez-Hernandez, I. E., Lin, X., Chang, Y. B., Rahman, A., Edwards, D. J. & Krakauer, J. W. The reliability of repeated TMS measures in older adults and in patients with subacute and chronic stroke. *Front Cell Neurosci.* (2015) **9**, 335.

2	Gertz, H., Lingnau, A. & Fiehler, K. Decoding Movement Goals from the Fronto-Parietal Reach Network. *Front Hum Neurosci.* (2017) **11**, 84.

3	Calhoun, V. D., Adali, T., Pearlson, G. D. & Pekar, J. J. A method for making group inferences from functional MRI data using independent component analysis. *Hum Brain Mapp.* (2001) **14**, 140-151.

4	Arbabshirani, M. R., Havlicek, M., Kiehl, K. A., Pearlson, G. D. & Calhoun, V. D. Functional network connectivity during rest and task conditions: a comparative study. *Hum Brain Mapp.* (2013) **34**, 2959-2971.

5	Rosazza, C. & Minati, L. Resting-state brain networks: literature review and clinical applications. *Neurol Sci.* (2011) **32**, 773-785.

6	Uddin, L. Q., Kelly, A. M., Biswal, B. B., Castellanos, F. X. & Milham, M. P. Functional connectivity of default mode network components: correlation, anticorrelation, and causality. *Hum Brain Mapp.* (2009) **30**, 625-637.

7	Fang, X., Zhang, Y., Zhou, Y., Cheng, L., Li, J., Wang, Y., Friston, K. J. & Jiang, T. Resting-State Coupling between Core Regions within the Central-Executive and Salience Networks Contributes to Working Memory Performance. *Front Behav Neurosci.* (2016) **10**, 27.

8	Ma, L., Wang, B., Chen, X. & Xiong, J. Detecting functional connectivity in the resting brain: a comparison between ICA and CCA. *Magn Reson Imaging.* (2007) **25**, 47-56.
